# A Programmable Model for Exploring the Functional Logic of the *Drosophila* Antennal Lobe

**DOI:** 10.1101/2022.09.10.506216

**Authors:** Aurel A. Lazar, Mehmet Kerem Turkcan, Yiyin Zhou

**Affiliations:** Department of Electrical Engineering, Columbia University, New York, NY 10027, USA

## Abstract

Recent progress on connectomics has resulted in huge datasets of brain structures in single-synapse scale. This calls for the modeling of executable circuits to discover the functional logic of these neural circuits in different scales. To provide an approach to standardize analyses of neuropils with multiple-input-multi-output channels, we put forward a programmable model focusing on the antennal lobe (AL), a circuit at the periphery of the olfactory system with nonlinear computation. We present an approach for constructing and analyzing antennal lobe circuits using models of glomeruli and local feedback loops. We establish the composability of the connectivity of glomeruli with local neuron feedback loops by combining pairs of glomeruli, and then by combining all glomeruli together to characterize the I/O of the entire AL. We thus provide a methodology for the quantitative characterization of the I/O of the AL as a function of feedback loop motifs.

## Executive Summary

### Characterizing the AL with Glomeruli Operating in Isolation

#### I/O Characterization of a Single Glomerulus

##### Objective

Single glomerulus parametrized by feedback loops. PN PSTH as a function of OSN PSTH (steady-state glomerulus I/O) for randomly generated number of synapses.

For a single glomerulus, we focus on the port connectivity patterns in Figure 1A. For each port connectivity pattern, we obtain the steady-state glomerulus I/Os (see Figure 1(B),(D),(E)). Here we used a generic glomerulus circuit model, whose parameters are a random subset of the number of postulated total number of synapses of each connected port. For each connected port, the number of synapses is randomly drawn between 0 and the maximum number of synapses observed in the Hemibrain dataset for that port.

**Figure 1:**
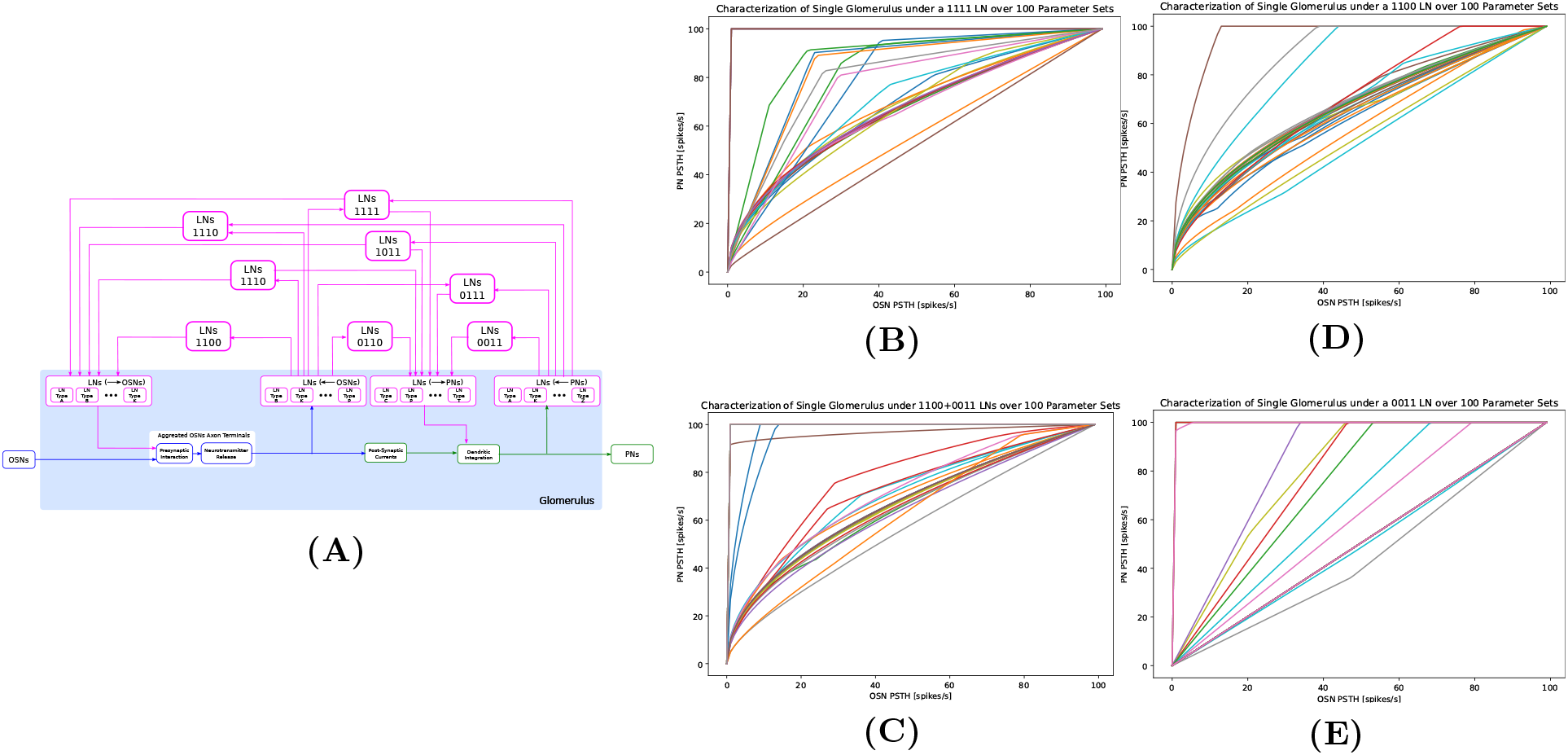
(A) Composability of the port connectivity of LNs within a single glomerulus. **(B-E)** Characterization of the effect of 3 port connectivity patterns and a simple port combination on the glomerulus I/O. (B) Port connectivity pattern 1111, (C) Port connectivity pattern 1100 and 0011, (D) Port connectivity pattern 1100, (E) Port connectivity pattern 0011. Each curve is the I/O result from one of the 100 parameter sets.

We also characterized the I/O of a glomerulus with a combination of port connectivity patterns 1100 and 0011 (Figure 1C), and noted that it can lead to very different I/O than the connectivity pattern 1111. This observation highlights the nonlinear nature of the I/O.

#### I/O Comparison Across Glomeruli

##### Objective

Single odorant at a given concentration level, I/O of all glomeruli (with number of synapses of the Hemibrain datasets) parametrized by the LN feedback loops.

We created a table of port connectivity patterns for all 50 glomeruli. Table 1 shows the number of neurons that has each of the patterns. Table 2 shows the total number of synapses for each port of each pattern in each glomerulus.

In Figure 2, we evaluate the I/O of the AL consisting of 50 glomeruli that operate in parallel. Each glomerulus is parametrized by a number of synapses for each connectivity pattern running independently in parallel. We pick an odorant with concentration fixed at 100ppm. We plot the PN responses of all glomeruli in Figure 2, one axis lists the glomeruli, the other axis lists the different LN pattern configurations. The effect of different feedback loops on the PN response for a given odorant at a particular concentration can be easily seen.

**Figure 2:**
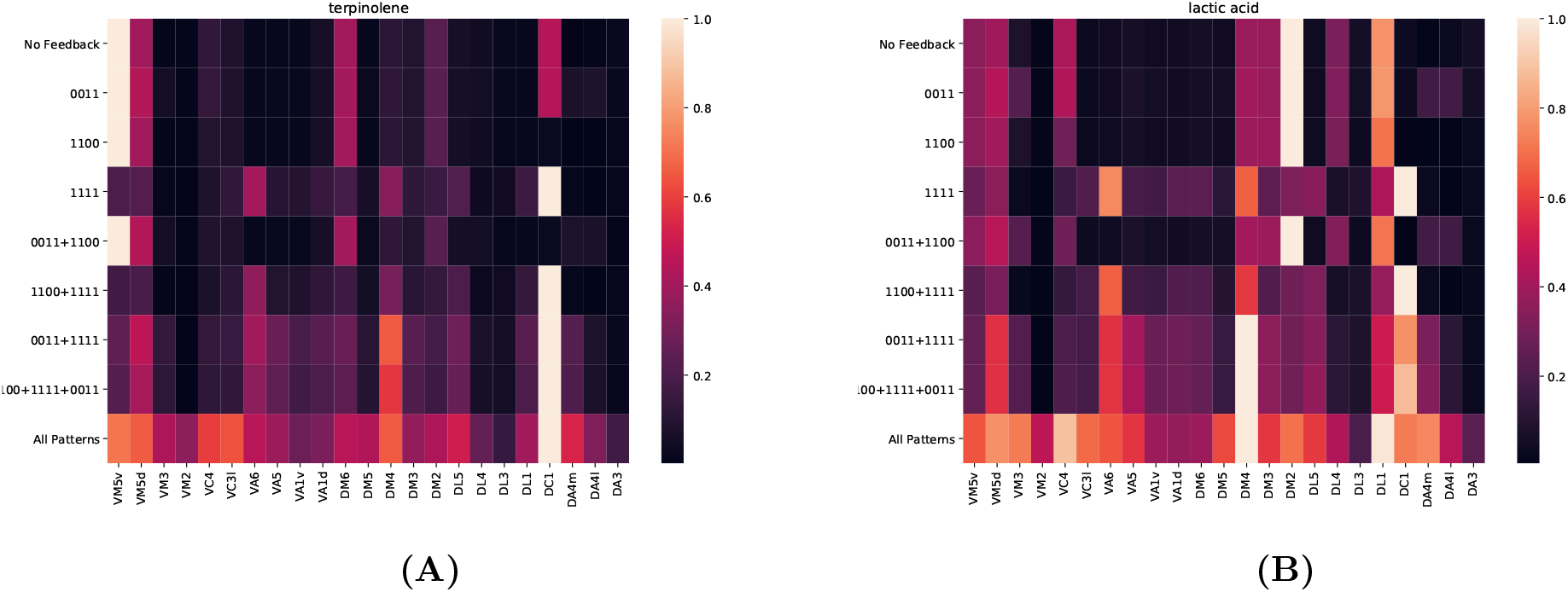
The responses of all 23 glomeruli run in isolation. Using an odorant **(A)** terpinolene and **(B)** lactic acid with 100ppm concentration as input, each entry indicates the response of PNs in a glomerulus (columns) and for each port connectivity pattern or combinations of patterns (rows) we consider.

### Characterization of Blocks of 2 Glomeruli with Feedback Motifs

From now on we assume that the local feedback in each of the glomeruli is intact (the existing local feedback will depend on the context). The motifs here will be connecting the glomeruli with existing local feedback loops. LN1 motif will interconnect all 1100 patterns, LN2 motif will interconnect all 0011 patterns, LN3 motif will interconnect all 1111 patterns. The goal is to compare the effect of motifs with over the baseline response of the two glomeruli circuit running in isolation.

#### Composition and I/O Characterization of a Block of Two Glomeruli

##### Objective

Block of two glomeruli, evaluate the change of PN PSTH of each glomerulus (in response to the OSN PSTHs at the input of each of the 2 glomeruli) due to each LN motif and with randomly generated number of synapses.

For a pair of glomeruli, we focus on the LN motif in Figure 3A, and characterize the I/O of the two glomeruli for a combination of individual motifs. Here we will fix a parameter set for glomerulus A and another parameter set for glomerulus B. Each glomerulus has the 1111, 1100, 0011 patterns in place. We execute the glomeruli in isolation to get the baseline response. First, we will only use the LN1 motif (connect all 1100 patterns), and show the difference in the response over baseline. Second, we will only use the LN2 motif (connect all 0011 patterns), and plot the difference. Third, LN3 motif. Fourth LN1+LN2 motif. So we will, in one figure (two subplots, one for uPN A, one for uPN B), have 4 surfaces of I/O change due to the different motifs. We show this in Figure 3B.

**Figure 3:**
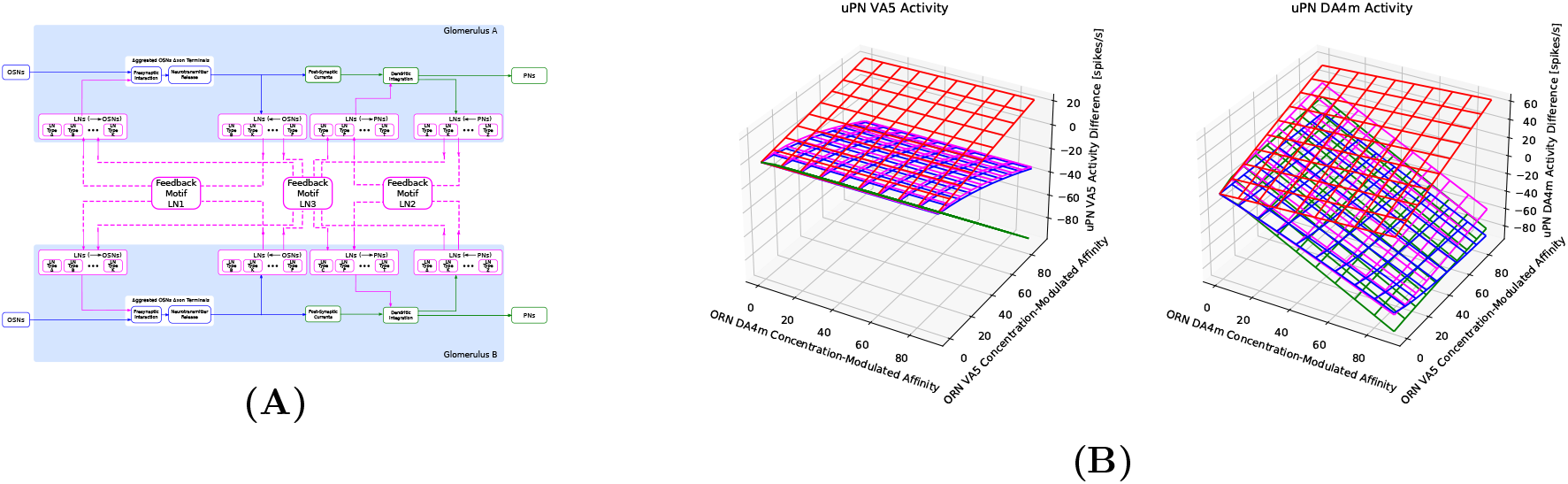
**(A)** LN feedback motifs for two glomeruli. **(B-C)** Characterization of I/O of a block of two glomeruli as a function of feedback motifs. **(B)** VA5 (left) and DA4m (right) PN responses when the two glomeruli run in isolation. **(C)** VA5(left) and DA4m (right) responses as a function of feedback motifs. (blue) The difference of PN responses to (B) when only feedback motif LN1 is used. (red) The difference of PN responses to (B) when only feedback motif LN2 is used. (green) The difference of PN responses to (B) when only feedback motif LN3 is used. (magenta) The difference of PN responses to (B) when both feedback motifs LN1 and LN2 are used.

#### Comparison Across Blocks of Two Glomeruli

##### Objective

Single odorant with constant concentration, evaluate the I/O of one glomerulus paired with all the other glomeruli. The I/O is parameterized by feedback motifs connecting glomerular pairs and the number of synapses is provided by the Hemibrain dataset.

Following Figure 2, we choose an odorant with a constant concentration. We obtain the baseline isolated glomerulus responses with their respective patterns 1100, 0011 and 1111. We then fix glomerulus A with the DL5 glomerulus, and pair it with a different glomerulus (D, DA1, DA2, etc.) in each run time execution. We plot how the DL5 glomerulus PN response (for the particular concentration and particular odorant) changes due to different combinations of feedback motifs and due to the pairing with the other glomeruli. The result is shown in Figure 4. The response of only one glomerulus is shown, and the response of the other glomeruli will appear, by symmetry, by repeating the above for all the glomeruli.

**Figure 4:**
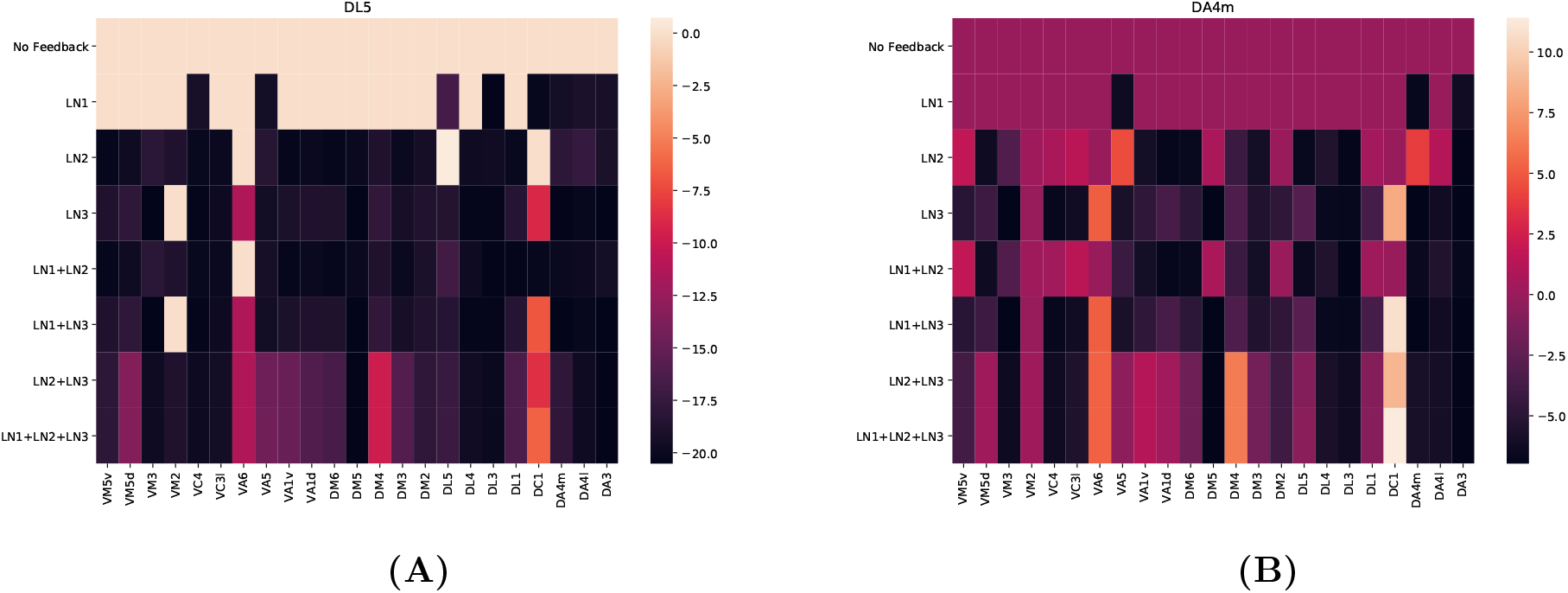
Evaluation of the I/O of a single glomerulus paired with all the other glomeruli. **(A)** DL5 glomerulus. **(B)** DA4m glomerulus. Each entry shows the difference of DL5 or DA4m PN response when paired with another glomerulus (indicated by the columns) over its response when isolated. Each row indicates a different combination of feedback motifs used to interconnect with the other glomerulus.

### Characterization of the AL as a Function of LN Motifs and Cell Types

#### I/O Characterization of AL Using Compositions of LN Feedback Motifs

##### Objective

Single odorant with a constant concentration level, evaluate I/O of all glomeruli parameterized by a global feedback motif and using the number of synapses for each port connectivity pattern of the Hemibrain dataset.

We extend Figure 3 to all glomeruli, and again characterize the AL with all glomeruli that now have LN motif 1, 2, 3 and combinations. For an odorant, we fix the concentration at 100ppm, and show the PN responses for all glomeruli in Figure 6.

#### I/O Characterization of AL with LN Cell Types

##### Objective

Single odorant with constant concentration, using Figure 5 and the Hemibrain dataset, evaluate the I/O of all glomeruli parameterized by LN types or individual LNs.

**Figure 5:**
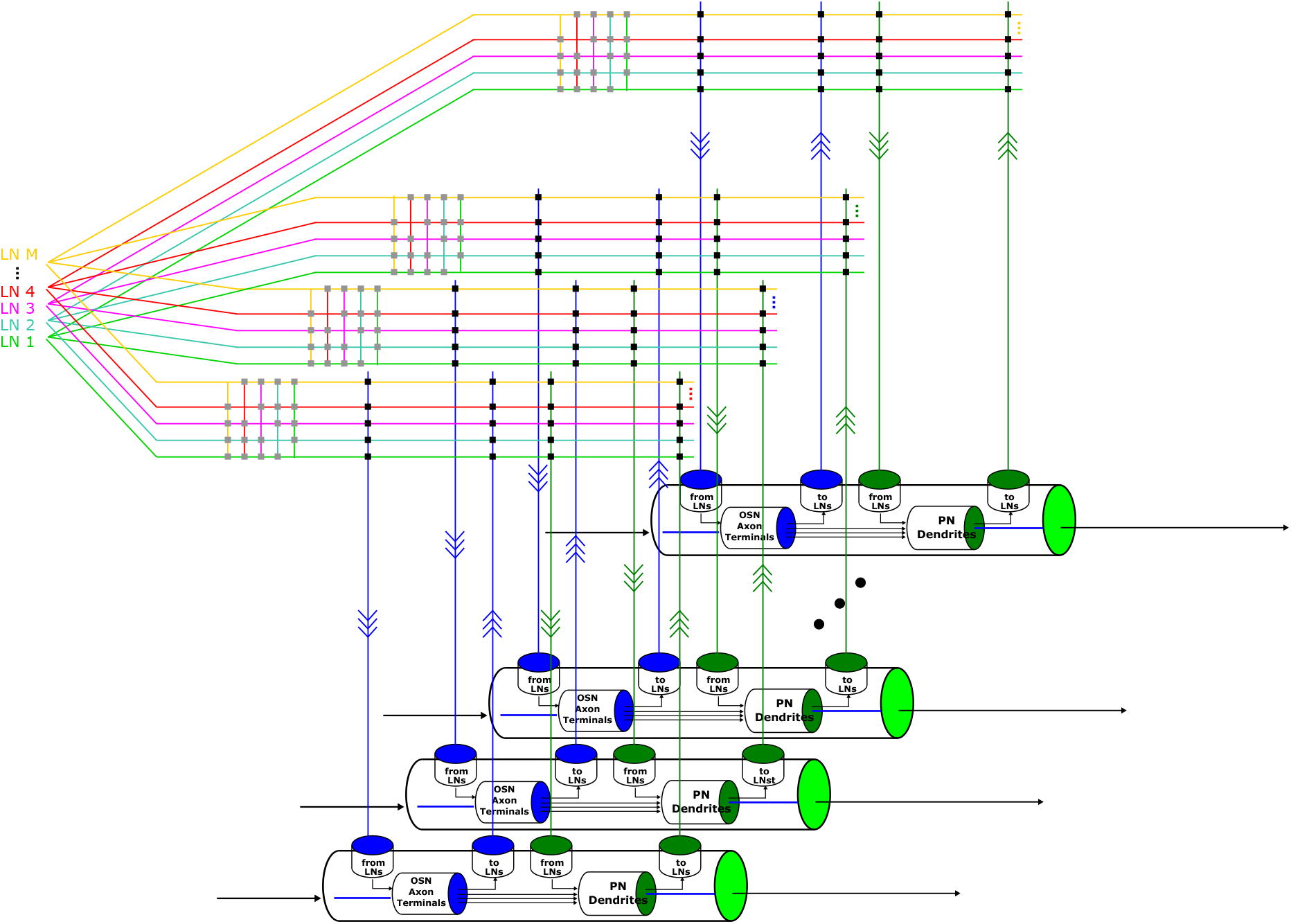
A circuit diagram modeling the entire AL circuit composed of the glomeruli and the massive number of feedback circuits.

**Figure 6:**
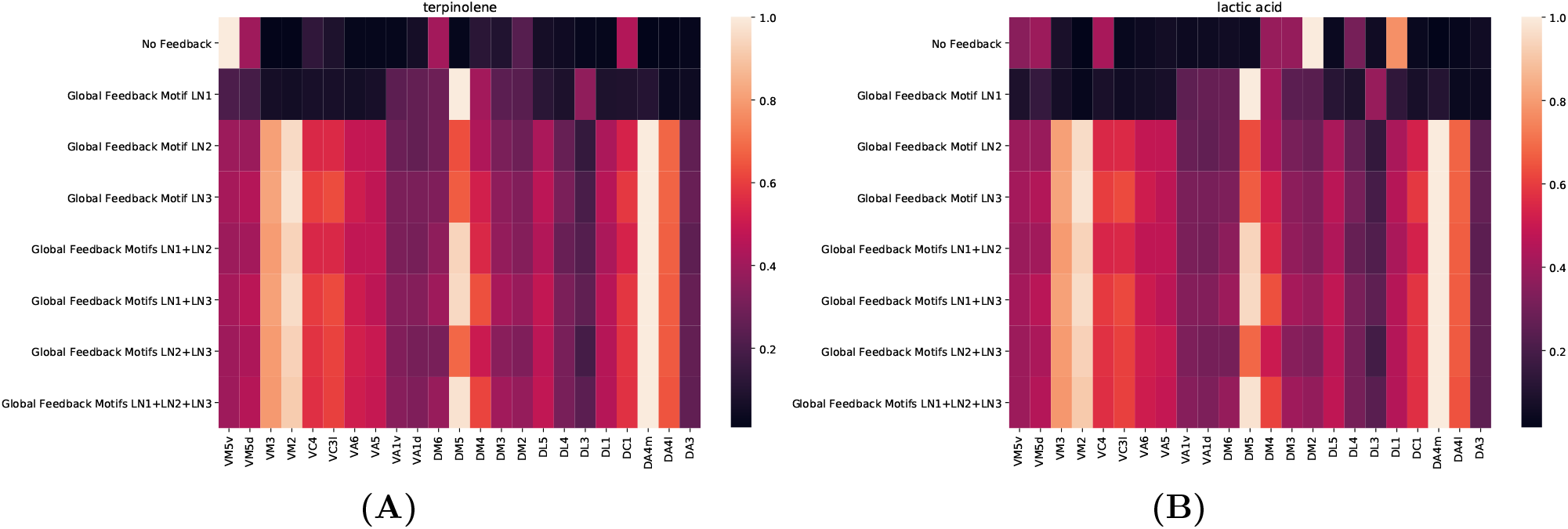
Comparison of the effect of individual LN motifs and combinations thereof on the I/O of all glomeruli, given a single odorant at 100ppm concentration level. **(A)** terpinolene. **(B)** lactic acid

We explore the I/O of the AL with respect to a single LN or a type of LNs. Either we only add the LNs of the single type, or ablate them from the full AL circuit. We also characterize the differences of the effect of adding a single, two, three, etc. neurons of an LN type.

In Figure 7, we start with all glomeruli with all 15 port connectivity patterns for local feedback intact. We then add all LNs of an LN type, connecting the synapses that these LNs form with each glomerulus (a subset of instances of each connectivity pattern). We execute the circuit by presenting an odorant with constant concentration of 100ppm. We show the PN responses for all glomeruli as a result of adding each LN type.

**Figure 7:**
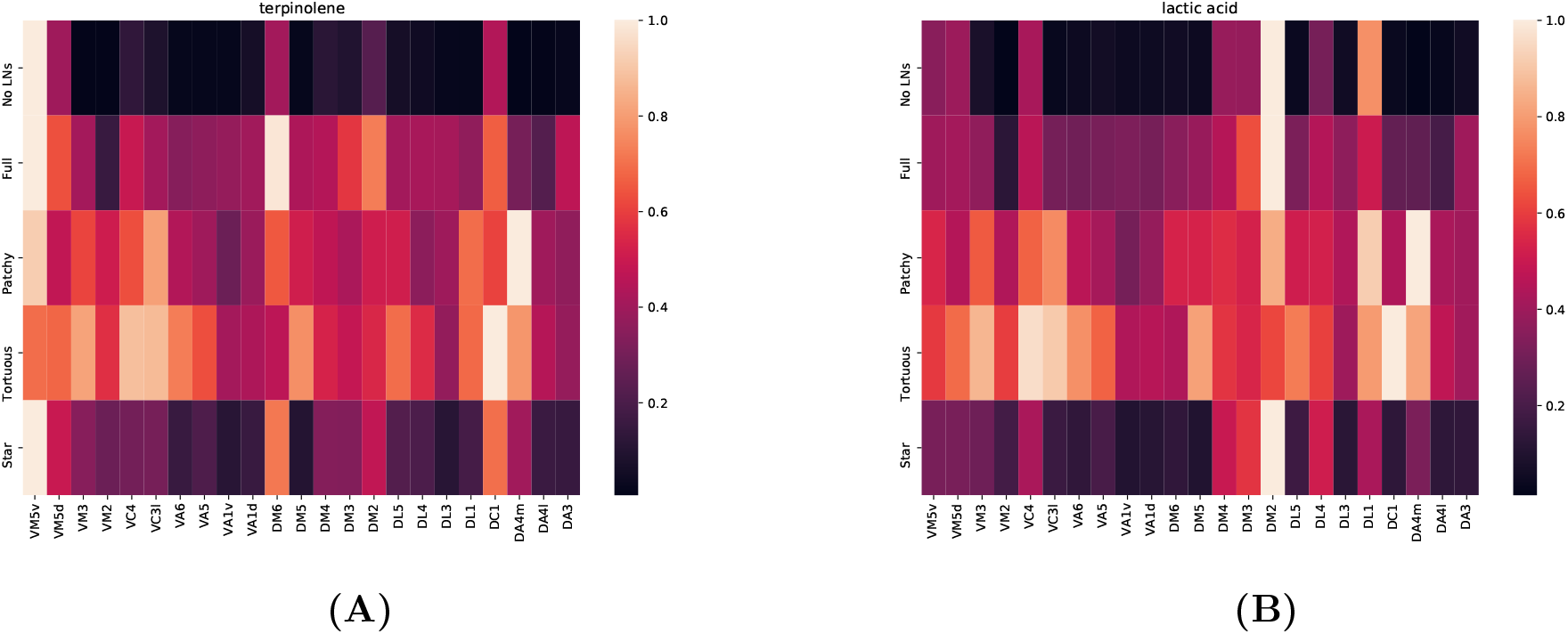
Comparison of the effect of individual LN types on the I/O of the entire AL when presented a single odorant. **(A)** terpinolene. **(B)** lactic acid.

In Figure 8, as in Figure 7, for a specific LN type we plot the PN PSTHs obtained after adding one by one LNs of the same type.

**Figure 8:**
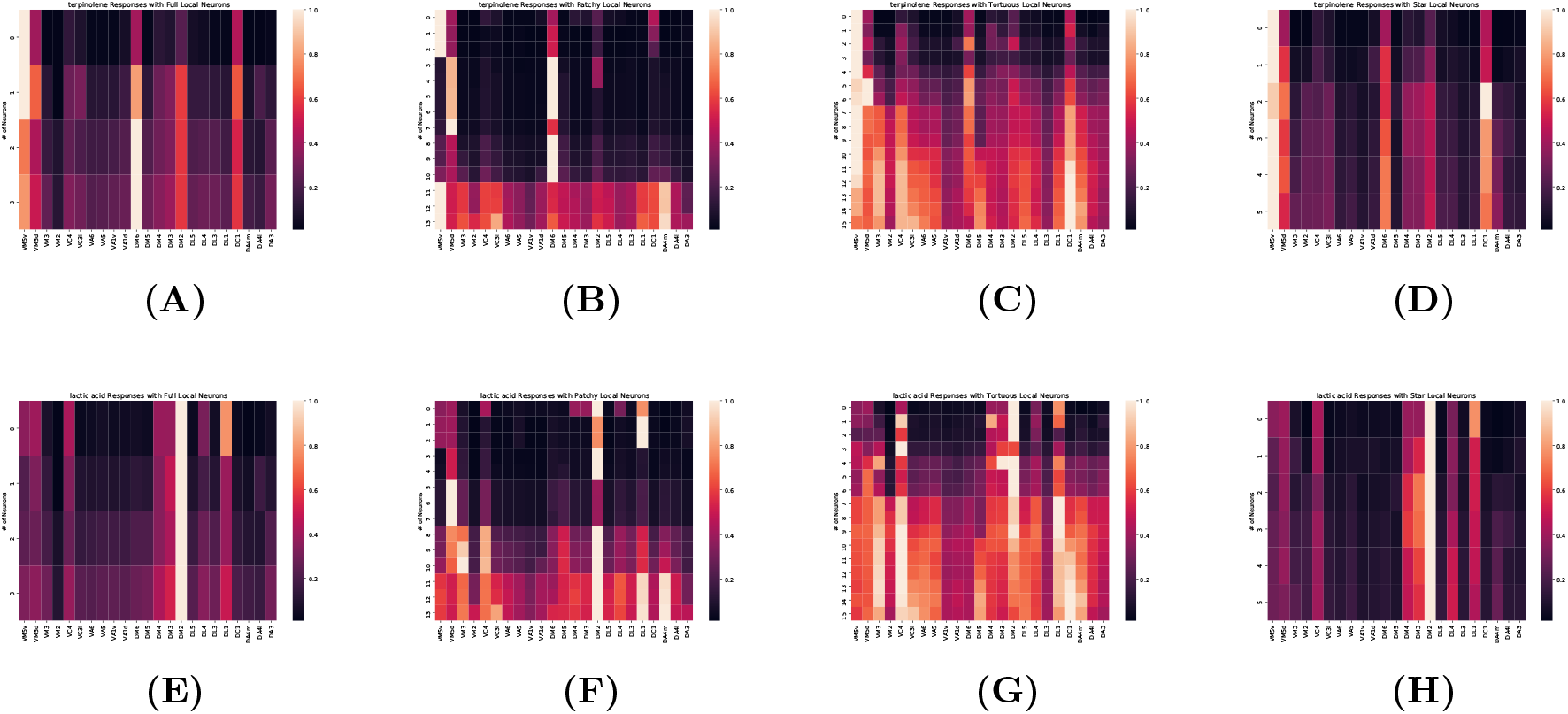
Comparison of the effect of individual LN neurons of an LN type on the I/O of the entire AL. **(A-D)** Responses of all glomeruli when presented terpinolene at 100ppm concentration level evaluated with adding LNs one by one of the (A) Full LNs, (B) Patchy LNs, (C) Tortuous LNs and (D) Star LNs. **(E-H)** Same as (A-D) but when presented lactic acid.

## Acknowledgments

The research reported here was supported by AFOSR under grant #FA9550-16-1-0410, DARPA under contract #HR0011-19-9-0035 and NSF under grant #2024607.

