## Supplementary material for "A Programmable Model for Exploring the Functional Logic of the *Drosophila* Antennal Lobe": Table 1: table1.html

|  | VP5 | VP3 | VP2 | VP1m | VP1l | VM7v | VM7d | VM5v | VM5d | VM4 | VM3 | VM2 | VM1 | VL2p | VL2a | VL1 | VC5 | VC4 | VC3m | VC3l | VC2 | VC1 | VA7m | VA7l | VA6 | VA5 | VA4 | VA3 | VA2 | VA1v | VA1d | V | DP1m | DP1l | DM6 | DM5 | DM4 | DM3 | DM2 | DM1 | DL5 | DL4 | DL3 | DL2v | DL2d | DL1 | DC4 | DC3 | DC2 | DC1 | DA4m | DA4l | DA3 | DA2 | DA1 | D |
| --- | --- | --- | --- | --- | --- | --- | --- | --- | --- | --- | --- | --- | --- | --- | --- | --- | --- | --- | --- | --- | --- | --- | --- | --- | --- | --- | --- | --- | --- | --- | --- | --- | --- | --- | --- | --- | --- | --- | --- | --- | --- | --- | --- | --- | --- | --- | --- | --- | --- | --- | --- | --- | --- | --- | --- | --- |
| 0011 | 30 | 40 | 93 | 66 | 62 | 12 | 9 | 5 | 20 | 13 | 29 | 8 | 23 | 4 | 2 | 26 | 8 | 9 | 13 | 10 | 11 | 4 | 6 | 23 | 1 | 23 | 12 | 7 | 2 | 17 | 11 | 10 | 6 | 10 | 10 | 17 | 14 | 11 | 8 | 3 | 9 | 14 | 7 | 3 | 13 | 13 | 6 | 3 | 8 | 3 | 39 | 33 | 6 | 8 | 6 | 32 |
| 1100 | 0 | 0 | 0 | 0 | 0 | 0 | 0 | 0 | 0 | 0 | 0 | 0 | 1 | 0 | 3 | 2 | 0 | 1 | 0 | 0 | 0 | 3 | 3 | 1 | 1 | 1 | 0 | 2 | 1 | 0 | 0 | 0 | 14 | 0 | 0 | 0 | 0 | 0 | 1 | 4 | 2 | 0 | 2 | 1 | 0 | 1 | 4 | 0 | 0 | 8 | 2 | 1 | 3 | 2 | 0 | 0 |
| 1111 | 0 | 0 | 0 | 0 | 0 | 35 | 49 | 44 | 46 | 25 | 10 | 0 | 23 | 54 | 55 | 45 | 51 | 32 | 31 | 43 | 33 | 42 | 44 | 24 | 56 | 39 | 25 | 36 | 46 | 46 | 45 | 51 | 60 | 49 | 52 | 18 | 58 | 48 | 52 | 75 | 52 | 26 | 28 | 50 | 38 | 44 | 66 | 49 | 57 | 59 | 12 | 4 | 17 | 36 | 46 | 72 |
| 0001 | 10 | 31 | 20 | 17 | 19 | 10 | 3 | 2 | 1 | 6 | 8 | 9 | 3 | 3 | 2 | 22 | 11 | 5 | 7 | 7 | 2 | 5 | 7 | 19 | 6 | 19 | 9 | 8 | 2 | 9 | 8 | 5 | 1 | 4 | 16 | 11 | 13 | 7 | 2 | 0 | 12 | 10 | 10 | 2 | 3 | 11 | 5 | 3 | 1 | 1 | 17 | 17 | 5 | 13 | 10 | 24 |
| 0010 | 55 | 48 | 36 | 21 | 18 | 16 | 15 | 4 | 14 | 12 | 4 | 14 | 10 | 17 | 11 | 34 | 17 | 12 | 18 | 10 | 5 | 9 | 10 | 12 | 11 | 14 | 11 | 7 | 0 | 16 | 11 | 19 | 7 | 8 | 8 | 12 | 25 | 19 | 18 | 17 | 7 | 18 | 5 | 4 | 12 | 21 | 12 | 12 | 9 | 10 | 20 | 17 | 6 | 4 | 10 | 18 |
| 0100 | 0 | 0 | 0 | 0 | 0 | 6 | 2 | 12 | 2 | 9 | 4 | 1 | 6 | 11 | 5 | 11 | 8 | 2 | 2 | 3 | 14 | 8 | 4 | 0 | 10 | 1 | 0 | 4 | 3 | 4 | 3 | 6 | 10 | 7 | 0 | 1 | 5 | 0 | 5 | 14 | 4 | 3 | 2 | 5 | 3 | 4 | 20 | 8 | 4 | 12 | 4 | 0 | 5 | 10 | 1 | 7 |
| 1000 | 0 | 0 | 0 | 0 | 0 | 0 | 1 | 1 | 0 | 1 | 0 | 1 | 0 | 1 | 2 | 0 | 2 | 0 | 0 | 0 | 1 | 1 | 0 | 0 | 0 | 0 | 1 | 0 | 0 | 0 | 2 | 1 | 1 | 2 | 0 | 1 | 2 | 1 | 1 | 1 | 0 | 0 | 1 | 3 | 0 | 1 | 1 | 2 | 1 | 2 | 0 | 0 | 1 | 0 | 0 | 0 |
| 1010 | 0 | 0 | 0 | 0 | 0 | 0 | 1 | 1 | 0 | 1 | 0 | 3 | 0 | 0 | 0 | 0 | 0 | 0 | 0 | 0 | 0 | 0 | 0 | 1 | 1 | 0 | 0 | 0 | 0 | 1 | 0 | 0 | 1 | 0 | 0 | 0 | 1 | 1 | 1 | 3 | 0 | 0 | 1 | 0 | 0 | 0 | 2 | 1 | 1 | 2 | 1 | 2 | 0 | 0 | 1 | 0 |
| 0101 | 0 | 0 | 0 | 0 | 0 | 2 | 5 | 4 | 3 | 3 | 0 | 1 | 0 | 9 | 6 | 16 | 7 | 3 | 3 | 1 | 5 | 4 | 3 | 4 | 7 | 6 | 5 | 6 | 3 | 4 | 8 | 6 | 4 | 9 | 2 | 1 | 4 | 2 | 3 | 4 | 3 | 3 | 2 | 1 | 5 | 5 | 3 | 6 | 8 | 8 | 3 | 2 | 1 | 5 | 3 | 8 |
| 0110 | 0 | 0 | 0 | 0 | 0 | 3 | 4 | 0 | 5 | 2 | 2 | 1 | 3 | 7 | 3 | 0 | 6 | 3 | 5 | 8 | 4 | 6 | 5 | 0 | 9 | 3 | 4 | 4 | 5 | 1 | 0 | 5 | 5 | 7 | 3 | 2 | 11 | 3 | 2 | 7 | 6 | 1 | 0 | 5 | 1 | 2 | 8 | 11 | 12 | 4 | 0 | 1 | 4 | 3 | 1 | 3 |
| 1001 | 0 | 0 | 0 | 0 | 0 | 0 | 0 | 0 | 1 | 0 | 0 | 0 | 1 | 0 | 1 | 0 | 0 | 0 | 0 | 0 | 1 | 0 | 0 | 0 | 0 | 0 | 0 | 0 | 0 | 0 | 0 | 0 | 0 | 0 | 0 | 0 | 0 | 0 | 2 | 1 | 0 | 1 | 0 | 1 | 0 | 0 | 0 | 0 | 0 | 0 | 0 | 1 | 0 | 1 | 0 | 1 |
| 0111 | 0 | 0 | 0 | 0 | 0 | 26 | 19 | 10 | 11 | 11 | 6 | 0 | 9 | 13 | 6 | 16 | 11 | 22 | 22 | 20 | 23 | 19 | 7 | 27 | 13 | 12 | 17 | 23 | 10 | 6 | 2 | 6 | 12 | 8 | 12 | 7 | 10 | 12 | 22 | 8 | 15 | 8 | 3 | 15 | 19 | 15 | 13 | 10 | 15 | 8 | 10 | 1 | 9 | 10 | 7 | 7 |
| 1011 | 0 | 0 | 0 | 0 | 0 | 3 | 6 | 1 | 2 | 2 | 4 | 1 | 1 | 4 | 0 | 1 | 1 | 2 | 2 | 5 | 1 | 7 | 4 | 3 | 0 | 2 | 3 | 1 | 5 | 3 | 4 | 0 | 2 | 4 | 9 | 1 | 3 | 3 | 1 | 0 | 4 | 4 | 3 | 1 | 1 | 2 | 0 | 3 | 2 | 1 | 4 | 4 | 2 | 1 | 6 | 3 |
| 1101 | 0 | 0 | 0 | 0 | 0 | 1 | 1 | 1 | 1 | 0 | 0 | 0 | 0 | 0 | 0 | 0 | 1 | 0 | 1 | 0 | 2 | 1 | 0 | 0 | 0 | 1 | 0 | 0 | 0 | 0 | 1 | 1 | 0 | 0 | 0 | 0 | 1 | 0 | 1 | 1 | 1 | 0 | 0 | 2 | 1 | 0 | 0 | 2 | 0 | 5 | 0 | 0 | 0 | 1 | 0 | 2 |
| 1110 | 0 | 0 | 0 | 0 | 0 | 6 | 4 | 3 | 5 | 1 | 1 | 0 | 4 | 4 | 8 | 4 | 8 | 4 | 3 | 1 | 6 | 3 | 4 | 2 | 1 | 2 | 5 | 8 | 9 | 4 | 2 | 3 | 11 | 9 | 3 | 0 | 8 | 10 | 5 | 5 | 4 | 1 | 1 | 9 | 5 | 4 | 15 | 4 | 3 | 3 | 2 | 2 | 6 | 2 | 0 | 2 |
