## Supplementary material for "A Programmable Model for Exploring the Functional Logic of the *Drosophila* Antennal Lobe": Table 2: table2.html

|  | VP5 | VP3 | VP2 | VP1m | VP1l | VM7v | VM7d | VM5v | VM5d | VM4 | VM3 | VM2 | VM1 | VL2p | VL2a | VL1 | VC5 | VC4 | VC3m | VC3l | VC2 | VC1 | VA7m | VA7l | VA6 | VA5 | VA4 | VA3 | VA2 | VA1v | VA1d | V | DP1m | DP1l | DM6 | DM5 | DM4 | DM3 | DM2 | DM1 | DL5 | DL4 | DL3 | DL2v | DL2d | DL1 | DC4 | DC3 | DC2 | DC1 | DA4m | DA4l | DA3 | DA2 | DA1 | D |
| --- | --- | --- | --- | --- | --- | --- | --- | --- | --- | --- | --- | --- | --- | --- | --- | --- | --- | --- | --- | --- | --- | --- | --- | --- | --- | --- | --- | --- | --- | --- | --- | --- | --- | --- | --- | --- | --- | --- | --- | --- | --- | --- | --- | --- | --- | --- | --- | --- | --- | --- | --- | --- | --- | --- | --- | --- |
| 0011 | [0, 0, 603, 229] | [0, 0, 328, 371] | [0, 0, 3015, 2902] | [0, 0, 1829, 3405] | [0, 0, 1433, 1355] | [0, 0, 64, 74] | [0, 0, 113, 98] | [0, 0, 21, 35] | [0, 0, 165, 194] | [0, 0, 126, 183] | [0, 0, 198, 282] | [0, 0, 15, 13] | [0, 0, 103, 182] | [0, 0, 14, 11] | [0, 0, 15, 32] | [0, 0, 316, 134] | [0, 0, 99, 65] | [0, 0, 158, 114] | [0, 0, 99, 134] | [0, 0, 40, 37] | [0, 0, 108, 246] | [0, 0, 38, 65] | [0, 0, 38, 56] | [0, 0, 202, 328] | [0, 0, 1, 1] | [0, 0, 236, 362] | [0, 0, 119, 213] | [0, 0, 77, 96] | [0, 0, 3, 12] | [0, 0, 147, 80] | [0, 0, 101, 54] | [0, 0, 78, 113] | [0, 0, 37, 14] | [0, 0, 80, 82] | [0, 0, 34, 50] | [0, 0, 40, 69] | [0, 0, 294, 292] | [0, 0, 95, 142] | [0, 0, 51, 31] | [0, 0, 25, 18] | [0, 0, 55, 57] | [0, 0, 81, 169] | [0, 0, 23, 30] | [0, 0, 34, 19] | [0, 0, 260, 334] | [0, 0, 98, 83] | [0, 0, 20, 10] | [0, 0, 10, 18] | [0, 0, 135, 167] | [0, 0, 14, 5] | [0, 0, 202, 392] | [0, 0, 173, 199] | [0, 0, 23, 26] | [0, 0, 19, 59] | [0, 0, 27, 23] | [0, 0, 224, 196] |
| 1100 | [0, 0, 0, 0] | [0, 0, 0, 0] | [0, 0, 0, 0] | [0, 0, 0, 0] | [0, 0, 0, 0] | [0, 0, 0, 0] | [0, 0, 0, 0] | [0, 0, 0, 0] | [0, 0, 0, 0] | [0, 0, 0, 0] | [0, 0, 0, 0] | [0, 0, 0, 0] | [1, 3, 0, 0] | [0, 0, 0, 0] | [8, 6, 0, 0] | [16, 31, 0, 0] | [0, 0, 0, 0] | [7, 18, 0, 0] | [0, 0, 0, 0] | [0, 0, 0, 0] | [0, 0, 0, 0] | [3, 4, 0, 0] | [3, 20, 0, 0] | [1, 1, 0, 0] | [13, 39, 0, 0] | [1, 7, 0, 0] | [0, 0, 0, 0] | [5, 27, 0, 0] | [1, 67, 0, 0] | [0, 0, 0, 0] | [0, 0, 0, 0] | [0, 0, 0, 0] | [2185, 1975, 0, 0] | [0, 0, 0, 0] | [0, 0, 0, 0] | [0, 0, 0, 0] | [0, 0, 0, 0] | [0, 0, 0, 0] | [1, 1, 0, 0] | [5, 12, 0, 0] | [3, 14, 0, 0] | [0, 0, 0, 0] | [4, 16, 0, 0] | [1, 1, 0, 0] | [0, 0, 0, 0] | [1, 1, 0, 0] | [16, 11, 0, 0] | [0, 0, 0, 0] | [0, 0, 0, 0] | [199, 326, 0, 0] | [6, 5, 0, 0] | [5, 3, 0, 0] | [20, 41, 0, 0] | [8, 27, 0, 0] | [0, 0, 0, 0] | [0, 0, 0, 0] |
| 1111 | [0, 0, 0, 0] | [0, 0, 0, 0] | [0, 0, 0, 0] | [0, 0, 0, 0] | [0, 0, 0, 0] | [569, 669, 527, 529] | [1052, 1407, 972, 1194] | [1295, 1928, 647, 982] | [1791, 3223, 1266, 1477] | [393, 796, 314, 625] | [95, 29, 89, 139] | [0, 0, 0, 0] | [201, 368, 217, 401] | [3064, 2789, 2256, 2421] | [2600, 3798, 2007, 2339] | [2913, 3134, 1444, 2048] | [2067, 3519, 1570, 1753] | [1270, 1374, 863, 831] | [529, 802, 744, 919] | [1507, 1630, 1155, 1170] | [976, 1233, 684, 1108] | [896, 1759, 1125, 1782] | [929, 1179, 904, 1095] | [437, 674, 498, 606] | [1440, 3351, 1390, 1583] | [789, 1222, 974, 1088] | [646, 1095, 599, 780] | [2013, 1708, 953, 888] | [2018, 2363, 1000, 1583] | [1988, 2342, 1759, 1477] | [1833, 2610, 1570, 1946] | [2841, 5512, 2087, 2611] | [6786, 7645, 3494, 3569] | [2767, 3015, 2500, 2474] | [2827, 3756, 1160, 1633] | [217, 285, 121, 143] | [3892, 4369, 2052, 2618] | [1751, 2373, 952, 1165] | [2632, 2926, 1219, 1513] | [9093, 10768, 2316, 3692] | [2214, 3378, 1121, 1693] | [922, 1204, 315, 555] | [633, 1917, 771, 755] | [2050, 2059, 1596, 1488] | [1012, 1363, 1190, 1958] | [2078, 2629, 1558, 2020] | [2785, 5036, 940, 1078] | [1492, 2129, 1362, 1447] | [1661, 2550, 1564, 1754] | [2418, 3437, 1841, 2774] | [38, 39, 119, 204] | [30, 24, 8, 10] | [277, 397, 89, 132] | [375, 824, 421, 515] | [2715, 7326, 3315, 2900] | [2585, 3191, 3821, 4320] |
| 0001 | [0, 0, 0, 21] | [0, 0, 0, 162] | [0, 0, 0, 217] | [0, 0, 0, 40] | [0, 0, 0, 40] | [0, 0, 0, 18] | [0, 0, 0, 3] | [0, 0, 0, 2] | [0, 0, 0, 2] | [0, 0, 0, 10] | [0, 0, 0, 9] | [0, 0, 0, 10] | [0, 0, 0, 4] | [0, 0, 0, 19] | [0, 0, 0, 2] | [0, 0, 0, 136] | [0, 0, 0, 39] | [0, 0, 0, 8] | [0, 0, 0, 18] | [0, 0, 0, 9] | [0, 0, 0, 8] | [0, 0, 0, 10] | [0, 0, 0, 14] | [0, 0, 0, 81] | [0, 0, 0, 8] | [0, 0, 0, 47] | [0, 0, 0, 40] | [0, 0, 0, 18] | [0, 0, 0, 2] | [0, 0, 0, 9] | [0, 0, 0, 29] | [0, 0, 0, 7] | [0, 0, 0, 1] | [0, 0, 0, 13] | [0, 0, 0, 30] | [0, 0, 0, 20] | [0, 0, 0, 40] | [0, 0, 0, 21] | [0, 0, 0, 6] | [0, 0, 0, 0] | [0, 0, 0, 40] | [0, 0, 0, 41] | [0, 0, 0, 25] | [0, 0, 0, 2] | [0, 0, 0, 7] | [0, 0, 0, 34] | [0, 0, 0, 6] | [0, 0, 0, 4] | [0, 0, 0, 2] | [0, 0, 0, 1] | [0, 0, 0, 53] | [0, 0, 0, 53] | [0, 0, 0, 8] | [0, 0, 0, 48] | [0, 0, 0, 14] | [0, 0, 0, 157] |
| 0010 | [0, 0, 813, 0] | [0, 0, 529, 0] | [0, 0, 301, 0] | [0, 0, 129, 0] | [0, 0, 93, 0] | [0, 0, 31, 0] | [0, 0, 36, 0] | [0, 0, 5, 0] | [0, 0, 27, 0] | [0, 0, 16, 0] | [0, 0, 14, 0] | [0, 0, 22, 0] | [0, 0, 20, 0] | [0, 0, 62, 0] | [0, 0, 44, 0] | [0, 0, 149, 0] | [0, 0, 55, 0] | [0, 0, 31, 0] | [0, 0, 47, 0] | [0, 0, 12, 0] | [0, 0, 14, 0] | [0, 0, 13, 0] | [0, 0, 30, 0] | [0, 0, 21, 0] | [0, 0, 44, 0] | [0, 0, 50, 0] | [0, 0, 55, 0] | [0, 0, 28, 0] | [0, 0, 0, 0] | [0, 0, 60, 0] | [0, 0, 49, 0] | [0, 0, 78, 0] | [0, 0, 25, 0] | [0, 0, 30, 0] | [0, 0, 62, 0] | [0, 0, 15, 0] | [0, 0, 98, 0] | [0, 0, 53, 0] | [0, 0, 35, 0] | [0, 0, 46, 0] | [0, 0, 22, 0] | [0, 0, 48, 0] | [0, 0, 7, 0] | [0, 0, 8, 0] | [0, 0, 33, 0] | [0, 0, 121, 0] | [0, 0, 15, 0] | [0, 0, 48, 0] | [0, 0, 20, 0] | [0, 0, 27, 0] | [0, 0, 55, 0] | [0, 0, 63, 0] | [0, 0, 6, 0] | [0, 0, 6, 0] | [0, 0, 25, 0] | [0, 0, 124, 0] |
| 0100 | [0, 0, 0, 0] | [0, 0, 0, 0] | [0, 0, 0, 0] | [0, 0, 0, 0] | [0, 0, 0, 0] | [0, 17, 0, 0] | [0, 4, 0, 0] | [0, 21, 0, 0] | [0, 165, 0, 0] | [0, 45, 0, 0] | [0, 8, 0, 0] | [0, 1, 0, 0] | [0, 24, 0, 0] | [0, 408, 0, 0] | [0, 13, 0, 0] | [0, 251, 0, 0] | [0, 47, 0, 0] | [0, 2, 0, 0] | [0, 7, 0, 0] | [0, 8, 0, 0] | [0, 119, 0, 0] | [0, 28, 0, 0] | [0, 16, 0, 0] | [0, 0, 0, 0] | [0, 131, 0, 0] | [0, 3, 0, 0] | [0, 0, 0, 0] | [0, 22, 0, 0] | [0, 64, 0, 0] | [0, 15, 0, 0] | [0, 12, 0, 0] | [0, 34, 0, 0] | [0, 67, 0, 0] | [0, 59, 0, 0] | [0, 0, 0, 0] | [0, 5, 0, 0] | [0, 7, 0, 0] | [0, 0, 0, 0] | [0, 35, 0, 0] | [0, 100, 0, 0] | [0, 11, 0, 0] | [0, 7, 0, 0] | [0, 8, 0, 0] | [0, 79, 0, 0] | [0, 46, 0, 0] | [0, 45, 0, 0] | [0, 123, 0, 0] | [0, 33, 0, 0] | [0, 15, 0, 0] | [0, 113, 0, 0] | [0, 11, 0, 0] | [0, 0, 0, 0] | [0, 14, 0, 0] | [0, 33, 0, 0] | [0, 1, 0, 0] | [0, 39, 0, 0] |
| 1000 | [0, 0, 0, 0] | [0, 0, 0, 0] | [0, 0, 0, 0] | [0, 0, 0, 0] | [0, 0, 0, 0] | [0, 0, 0, 0] | [1, 0, 0, 0] | [1, 0, 0, 0] | [0, 0, 0, 0] | [1, 0, 0, 0] | [0, 0, 0, 0] | [1, 0, 0, 0] | [0, 0, 0, 0] | [1, 0, 0, 0] | [4, 0, 0, 0] | [0, 0, 0, 0] | [2, 0, 0, 0] | [0, 0, 0, 0] | [0, 0, 0, 0] | [0, 0, 0, 0] | [1, 0, 0, 0] | [1, 0, 0, 0] | [0, 0, 0, 0] | [0, 0, 0, 0] | [0, 0, 0, 0] | [0, 0, 0, 0] | [1, 0, 0, 0] | [0, 0, 0, 0] | [0, 0, 0, 0] | [0, 0, 0, 0] | [2, 0, 0, 0] | [1, 0, 0, 0] | [1, 0, 0, 0] | [3, 0, 0, 0] | [0, 0, 0, 0] | [2, 0, 0, 0] | [2, 0, 0, 0] | [1, 0, 0, 0] | [4, 0, 0, 0] | [1, 0, 0, 0] | [0, 0, 0, 0] | [0, 0, 0, 0] | [1, 0, 0, 0] | [4, 0, 0, 0] | [0, 0, 0, 0] | [2, 0, 0, 0] | [2, 0, 0, 0] | [3, 0, 0, 0] | [1, 0, 0, 0] | [2, 0, 0, 0] | [0, 0, 0, 0] | [0, 0, 0, 0] | [4, 0, 0, 0] | [0, 0, 0, 0] | [0, 0, 0, 0] | [0, 0, 0, 0] |
| 1010 | [0, 0, 0, 0] | [0, 0, 0, 0] | [0, 0, 0, 0] | [0, 0, 0, 0] | [0, 0, 0, 0] | [0, 0, 0, 0] | [1, 0, 1, 0] | [1, 0, 1, 0] | [0, 0, 0, 0] | [2, 0, 2, 0] | [0, 0, 0, 0] | [4, 0, 4, 0] | [0, 0, 0, 0] | [0, 0, 0, 0] | [0, 0, 0, 0] | [0, 0, 0, 0] | [0, 0, 0, 0] | [0, 0, 0, 0] | [0, 0, 0, 0] | [0, 0, 0, 0] | [0, 0, 0, 0] | [0, 0, 0, 0] | [0, 0, 0, 0] | [1, 0, 2, 0] | [1, 0, 1, 0] | [0, 0, 0, 0] | [0, 0, 0, 0] | [0, 0, 0, 0] | [0, 0, 0, 0] | [1, 0, 7, 0] | [0, 0, 0, 0] | [0, 0, 0, 0] | [1, 0, 4, 0] | [0, 0, 0, 0] | [0, 0, 0, 0] | [0, 0, 0, 0] | [1, 0, 2, 0] | [1, 0, 1, 0] | [2, 0, 5, 0] | [12, 0, 15, 0] | [0, 0, 0, 0] | [0, 0, 0, 0] | [13, 0, 1, 0] | [0, 0, 0, 0] | [0, 0, 0, 0] | [0, 0, 0, 0] | [2, 0, 4, 0] | [3, 0, 8, 0] | [1, 0, 12, 0] | [4, 0, 19, 0] | [2, 0, 2, 0] | [3, 0, 2, 0] | [0, 0, 0, 0] | [0, 0, 0, 0] | [1, 0, 1, 0] | [0, 0, 0, 0] |
| 0101 | [0, 0, 0, 0] | [0, 0, 0, 0] | [0, 0, 0, 0] | [0, 0, 0, 0] | [0, 0, 0, 0] | [0, 2, 0, 2] | [0, 22, 0, 32] | [0, 23, 0, 16] | [0, 8, 0, 16] | [0, 66, 0, 4] | [0, 0, 0, 0] | [0, 1, 0, 1] | [0, 0, 0, 0] | [0, 125, 0, 105] | [0, 366, 0, 22] | [0, 1970, 0, 38] | [0, 213, 0, 35] | [0, 15, 0, 11] | [0, 9, 0, 5] | [0, 3, 0, 4] | [0, 7, 0, 6] | [0, 22, 0, 40] | [0, 10, 0, 15] | [0, 46, 0, 18] | [0, 184, 0, 19] | [0, 33, 0, 63] | [0, 47, 0, 50] | [0, 277, 0, 31] | [0, 44, 0, 13] | [0, 72, 0, 15] | [0, 70, 0, 98] | [0, 84, 0, 33] | [0, 213, 0, 33] | [0, 361, 0, 48] | [0, 4, 0, 10] | [0, 1, 0, 6] | [0, 75, 0, 29] | [0, 12, 0, 5] | [0, 47, 0, 16] | [0, 41, 0, 5] | [0, 26, 0, 4] | [0, 5, 0, 7] | [0, 5, 0, 10] | [0, 5, 0, 4] | [0, 18, 0, 17] | [0, 27, 0, 36] | [0, 17, 0, 4] | [0, 28, 0, 9] | [0, 49, 0, 26] | [0, 63, 0, 25] | [0, 7, 0, 31] | [0, 12, 0, 4] | [0, 1, 0, 14] | [0, 28, 0, 20] | [0, 41, 0, 7] | [0, 37, 0, 22] |
| 0110 | [0, 0, 0, 0] | [0, 0, 0, 0] | [0, 0, 0, 0] | [0, 0, 0, 0] | [0, 0, 0, 0] | [0, 8, 33, 0] | [0, 7, 15, 0] | [0, 0, 0, 0] | [0, 9, 61, 0] | [0, 7, 3, 0] | [0, 11, 8, 0] | [0, 2, 3, 0] | [0, 92, 4, 0] | [0, 22, 39, 0] | [0, 8, 23, 0] | [0, 0, 0, 0] | [0, 12, 23, 0] | [0, 11, 22, 0] | [0, 28, 28, 0] | [0, 19, 21, 0] | [0, 9, 13, 0] | [0, 41, 22, 0] | [0, 34, 63, 0] | [0, 0, 0, 0] | [0, 150, 33, 0] | [0, 10, 6, 0] | [0, 46, 10, 0] | [0, 60, 26, 0] | [0, 25, 30, 0] | [0, 1, 7, 0] | [0, 0, 0, 0] | [0, 702, 14, 0] | [0, 118, 20, 0] | [0, 279, 33, 0] | [0, 15, 12, 0] | [0, 17, 7, 0] | [0, 128, 28, 0] | [0, 7, 15, 0] | [0, 5, 24, 0] | [0, 20, 61, 0] | [0, 14, 15, 0] | [0, 1, 2, 0] | [0, 0, 0, 0] | [0, 65, 32, 0] | [0, 3, 20, 0] | [0, 34, 4, 0] | [0, 85, 16, 0] | [0, 50, 79, 0] | [0, 102, 104, 0] | [0, 15, 14, 0] | [0, 0, 0, 0] | [0, 4, 1, 0] | [0, 76, 21, 0] | [0, 16, 11, 0] | [0, 2, 2, 0] | [0, 4, 34, 0] |
| 1001 | [0, 0, 0, 0] | [0, 0, 0, 0] | [0, 0, 0, 0] | [0, 0, 0, 0] | [0, 0, 0, 0] | [0, 0, 0, 0] | [0, 0, 0, 0] | [0, 0, 0, 0] | [2, 0, 0, 1] | [0, 0, 0, 0] | [0, 0, 0, 0] | [0, 0, 0, 0] | [1, 0, 0, 1] | [0, 0, 0, 0] | [1, 0, 0, 1] | [0, 0, 0, 0] | [0, 0, 0, 0] | [0, 0, 0, 0] | [0, 0, 0, 0] | [0, 0, 0, 0] | [1, 0, 0, 1] | [0, 0, 0, 0] | [0, 0, 0, 0] | [0, 0, 0, 0] | [0, 0, 0, 0] | [0, 0, 0, 0] | [0, 0, 0, 0] | [0, 0, 0, 0] | [0, 0, 0, 0] | [0, 0, 0, 0] | [0, 0, 0, 0] | [0, 0, 0, 0] | [0, 0, 0, 0] | [0, 0, 0, 0] | [0, 0, 0, 0] | [0, 0, 0, 0] | [0, 0, 0, 0] | [0, 0, 0, 0] | [2, 0, 0, 2] | [4, 0, 0, 2] | [0, 0, 0, 0] | [2, 0, 0, 3] | [0, 0, 0, 0] | [1, 0, 0, 1] | [0, 0, 0, 0] | [0, 0, 0, 0] | [0, 0, 0, 0] | [0, 0, 0, 0] | [0, 0, 0, 0] | [0, 0, 0, 0] | [0, 0, 0, 0] | [1, 0, 0, 3] | [0, 0, 0, 0] | [4, 0, 0, 35] | [0, 0, 0, 0] | [1, 0, 0, 2] |
| 0111 | [0, 0, 0, 0] | [0, 0, 0, 0] | [0, 0, 0, 0] | [0, 0, 0, 0] | [0, 0, 0, 0] | [0, 262, 237, 365] | [0, 145, 258, 258] | [0, 68, 172, 200] | [0, 63, 148, 99] | [0, 64, 220, 274] | [0, 17, 47, 77] | [0, 0, 0, 0] | [0, 19, 74, 136] | [0, 492, 147, 339] | [0, 111, 134, 228] | [0, 511, 122, 375] | [0, 1003, 184, 370] | [0, 131, 346, 327] | [0, 132, 339, 502] | [0, 169, 471, 515] | [0, 436, 343, 606] | [0, 313, 412, 494] | [0, 40, 58, 78] | [0, 216, 303, 608] | [0, 311, 89, 181] | [0, 168, 120, 215] | [0, 236, 189, 366] | [0, 284, 658, 691] | [0, 58, 104, 151] | [0, 64, 146, 201] | [0, 15, 18, 75] | [0, 791, 99, 87] | [0, 124, 266, 380] | [0, 231, 311, 412] | [0, 46, 76, 98] | [0, 46, 44, 71] | [0, 140, 133, 164] | [0, 90, 119, 202] | [0, 180, 357, 407] | [0, 150, 82, 92] | [0, 176, 203, 362] | [0, 393, 67, 191] | [0, 20, 34, 71] | [0, 182, 383, 485] | [0, 157, 553, 705] | [0, 55, 238, 331] | [0, 147, 128, 69] | [0, 118, 122, 214] | [0, 237, 200, 326] | [0, 162, 107, 158] | [0, 23, 93, 176] | [0, 4, 6, 12] | [0, 221, 35, 134] | [0, 56, 109, 92] | [0, 25, 134, 53] | [0, 22, 140, 115] |
| 1011 | [0, 0, 0, 0] | [0, 0, 0, 0] | [0, 0, 0, 0] | [0, 0, 0, 0] | [0, 0, 0, 0] | [5, 0, 25, 35] | [9, 0, 91, 109] | [1, 0, 9, 24] | [2, 0, 43, 50] | [4, 0, 47, 45] | [8, 0, 26, 40] | [1, 0, 3, 3] | [2, 0, 1, 1] | [21, 0, 144, 162] | [0, 0, 0, 0] | [21, 0, 2, 1] | [2, 0, 23, 35] | [5, 0, 57, 46] | [2, 0, 17, 30] | [6, 0, 113, 84] | [1, 0, 23, 37] | [10, 0, 137, 314] | [7, 0, 87, 123] | [17, 0, 50, 76] | [0, 0, 0, 0] | [2, 0, 33, 47] | [5, 0, 53, 93] | [1, 0, 33, 28] | [6, 0, 107, 111] | [16, 0, 102, 139] | [9, 0, 64, 76] | [0, 0, 0, 0] | [6, 0, 115, 95] | [8, 0, 124, 83] | [17, 0, 103, 182] | [2, 0, 5, 11] | [3, 0, 76, 90] | [3, 0, 41, 49] | [3, 0, 24, 27] | [0, 0, 0, 0] | [7, 0, 63, 111] | [4, 0, 38, 56] | [4, 0, 11, 12] | [1, 0, 31, 41] | [1, 0, 41, 64] | [5, 0, 31, 96] | [0, 0, 0, 0] | [9, 0, 33, 33] | [3, 0, 64, 64] | [2, 0, 37, 44] | [9, 0, 25, 67] | [4, 0, 27, 15] | [3, 0, 12, 11] | [1, 0, 7, 1] | [18, 0, 143, 191] | [19, 0, 247, 265] |
| 1101 | [0, 0, 0, 0] | [0, 0, 0, 0] | [0, 0, 0, 0] | [0, 0, 0, 0] | [0, 0, 0, 0] | [1, 1, 0, 3] | [1, 3, 0, 1] | [1, 1, 0, 1] | [2, 1, 0, 1] | [0, 0, 0, 0] | [0, 0, 0, 0] | [0, 0, 0, 0] | [0, 0, 0, 0] | [0, 0, 0, 0] | [0, 0, 0, 0] | [0, 0, 0, 0] | [1, 18, 0, 4] | [0, 0, 0, 0] | [1, 20, 0, 1] | [0, 0, 0, 0] | [10, 3, 0, 4] | [8, 8, 0, 1] | [0, 0, 0, 0] | [0, 0, 0, 0] | [0, 0, 0, 0] | [12, 13, 0, 2] | [0, 0, 0, 0] | [0, 0, 0, 0] | [0, 0, 0, 0] | [0, 0, 0, 0] | [1, 8, 0, 8] | [1, 46, 0, 21] | [0, 0, 0, 0] | [0, 0, 0, 0] | [0, 0, 0, 0] | [0, 0, 0, 0] | [1, 70, 0, 1] | [0, 0, 0, 0] | [7, 4, 0, 2] | [1, 6, 0, 1] | [1, 17, 0, 3] | [0, 0, 0, 0] | [0, 0, 0, 0] | [12, 8, 0, 4] | [1, 2, 0, 1] | [0, 0, 0, 0] | [0, 0, 0, 0] | [6, 33, 0, 63] | [0, 0, 0, 0] | [5, 51, 0, 10] | [0, 0, 0, 0] | [0, 0, 0, 0] | [0, 0, 0, 0] | [43, 26, 0, 2] | [0, 0, 0, 0] | [2, 22, 0, 5] |
| 1110 | [0, 0, 0, 0] | [0, 0, 0, 0] | [0, 0, 0, 0] | [0, 0, 0, 0] | [0, 0, 0, 0] | [135, 324, 80, 0] | [46, 122, 62, 0] | [21, 111, 53, 0] | [26, 237, 96, 0] | [20, 33, 2, 0] | [6, 7, 11, 0] | [0, 0, 0, 0] | [44, 63, 27, 0] | [48, 149, 86, 0] | [89, 85, 72, 0] | [51, 243, 10, 0] | [464, 535, 77, 0] | [5, 15, 69, 0] | [18, 45, 23, 0] | [15, 72, 4, 0] | [170, 405, 78, 0] | [86, 231, 53, 0] | [10, 52, 37, 0] | [21, 85, 37, 0] | [85, 174, 14, 0] | [24, 63, 34, 0] | [140, 459, 85, 0] | [131, 272, 157, 0] | [142, 199, 92, 0] | [231, 312, 198, 0] | [46, 3, 45, 0] | [65, 103, 201, 0] | [494, 1201, 179, 0] | [277, 344, 171, 0] | [4, 22, 32, 0] | [0, 0, 0, 0] | [18, 364, 180, 0] | [477, 819, 163, 0] | [155, 394, 99, 0] | [11, 31, 43, 0] | [120, 424, 40, 0] | [1, 148, 9, 0] | [1, 1, 1, 0] | [316, 619, 239, 0] | [81, 182, 229, 0] | [27, 503, 138, 0] | [115, 288, 74, 0] | [19, 134, 50, 0] | [4, 69, 46, 0] | [9, 52, 37, 0] | [3, 4, 14, 0] | [9, 5, 10, 0] | [37, 82, 16, 0] | [6, 33, 19, 0] | [0, 0, 0, 0] | [26, 43, 20, 0] |
